# Decoding Duchenne muscular dystrophy transcriptome to single nuclei level reveals clinical-genetic correlations

**DOI:** 10.1101/2023.03.01.530728

**Authors:** Xavier Suárez-Calvet, Esther Fernández-Simón, Daniel Natera, Cristina Jou, Patricia Pinol-Jurado, Elisa Villalobos, Carlos Ortez, Alexandra Monceau, Marianela Schiava, José Verdu-Díaz, James Clark, Zoe Laidler, Priyanka Mehra, Rasya Gokul-Nath, Jorge Alonso-Perez, Chiara Marini-Bettolo, Giorgio Tasca, Volker Straub, Michela Guglieri, Andrés Nascimento, Jordi Diaz-Manera

## Abstract

The cellular and molecular consequences of lack of dystrophin in humans are only partially known, which is crucial for the development of new therapies aiming to slow or stop the progression Duchenne and Becker muscular dystrophies. We analyzed muscle biopsies of DMD patients and controls using single nuclei RNA sequencing (snRNAseq) and correlated the results with clinical data. DMD samples displayed an increase in regenerative fibers, satellite cells and fibro-adipogenic progenitor cells (FAPs) and a decrease in slow fibers and smooth muscle cells. Samples from patients with stable mild weakness were characterized by an increase in regenerative fibers, while those from patients with progressive weakness had fewer muscle fibers and increased FAPs. DMD muscle fibers displayed a strong regenerative signature, while DMD FAPs upregulated genes producing extracellular matrix and molecules involved in several signaling pathways. An analysis of intercellular communication profile identified FAPs as a key regulator of cell signaling in DMD samples. We show significant differences in the gene expression profiled of the different cell populations present in DMD muscle samples compared to controls.

## 1. Introduction

Duchenne muscular dystrophy (DMD) is a genetic disorder produced by mutations in the dystrophin gene and characterized by early onset progressive muscle weakness leading to irreversible severe disability^1, 2^. Treatment with corticosteroids is part of the standard for care as they delay disease progression, although unfortunately do not change substantially the natural history of the disease^1, 3, 4^. Several new therapies have been tested or are still under research in clinical trials using different strategies, including but not limited to cell therapy, gene therapy, anti-inflammatory, pro-regenerative and, antioxidant drugs^5^.

Dystrophin is localized in the cytoplasmic face of the muscle membrane linking the sarcomeric proteins to the sarcolemma and the extracellular matrix (ECM)^6^. Muscle fibers lacking dystrophin are injured during muscle contraction leading to continuous cycles of muscle regeneration that ultimately fails resulting in loss of muscle fibers and expansion of fat and fibrotic tissue. The process of muscle degeneration in DMD involves a complex interplay between muscle fibers, muscle resident cells such as satellite cells and fibroadipogenic progenitor cells (FAPs) and, circulating cells infiltrating the muscle such as macrophages and lymphocytes^7, 8^. Despite considerable progress in the understanding of the degenerative process in DMD, there is still a considerable lack of knowledge of what are the cellular and molecular consequences of the absence of dystrophin in humans^9^. Most of the studies performed in humans have analyzed muscle samples using bulk proteomics or RNA identifying dysregulated molecular pathways in DMD, but these studies are not able to characterize what cells are responsible of each pathway, or how cells interplay to each other during the process of muscle degeneration^10-13^.

Single cell RNA sequencing (scRNAseq) and single nuclei RNA sequencing (snRNAseq) allow the analysis of gene expression to the single cell/nuclei levels using small fragments of tissue^14^. This technology enables the identification of the cell populations present in the tissue of interest in healthy and disease conditions, the study of differentially expressed genes in each cell population compared to controls and, the identification of potential communications between cells easing the understanding of how the disease process is orchestrated and its dynamics along disease progression^15, 16^. In the case of DMD, both scRNAseq and snRNAseq offer a unique opportunity to study the changes in gene expression profile in different muscle cell populations using biopsies that were obtained for diagnosis and are stored in biobanks. SnRNAseq offers some advantages in the study of skeletal muscle biopsies compared to scRNAseq. First, snRNAseq allows to study the gene expression of myofibers nuclei, which represent more than 80% of the nuclei present in muscle and are irremediably lost if scRNAseq is performed, as this technology requires viable cell suspension for sequencing and muscle fibers, which spread from one tendon to the other, and are therefore not entire when muscle biopsies of patients are obtained^16-19^. Second, adipocytes fragility difficult their inclusion in scRNAseq studies with the risk of losing an important contributor to transcriptomic variability in the case of muscular dystrophies, such as DMD, while this difficulty is reduced if snRNAseq is used for the analysis^15^. Finally, scRNAseq does not perform well in frozen tissues, requiring fresh muscle, which is a clear limitation due to the reduced availability of muscle tissue from already diagnosed patients^20^. There have been some snRNAseq studies published son far using murine models of DMD that have provided valuable clues about the nature of the process of muscle degeneration, but the information coming from human samples is very limited ^21, 22^. Here, we have applied snRNAseq to muscle samples obtained from DMD patients that were biopsied between the age of two and four years old, at early stages of the disease before steroids were started. We have implemented a protocol that has allowed us to use a minimum amount of tissue, around 25 mg of frozen muscle and obtain approximately between 10 to 20 thousand nuclei for the analysis^16, 23^. We are comparing the gene expression profile to the single nuclei level of these samples with age and gender matched controls to understand what are the cellular and molecular consequences of the lack of dystrophin that influence the process of muscle degeneration in humans.

## 2. Results

### Patients and samples included

We performed snRNAseq on 7 muscle samples of patients with DMD and 5 muscle samples of age and gender matched controls. Table 1 summarizes the main demographic, genetic and clinical features of the patients included. Supplemental figure 1 show representative areas of the haematoxylin-eosin (HE) staining of the muscle samples included in this study.

**Table 1:** Clinical and genetic data of DMD patients and control included in the study. G: gender. Quad: quadriceps. NSAA: North Star Ambulatory Assessment. 6MWT: Six minutes walking test. 10MWT: 10 minutes walking test. TSUF: Time to stand up from floor. S: seconds. M: meters. Y: years.

| Code | Age | G. | Muscle | Mutation | NSAA<br>Baseline | 6MWT<br>baseline | 10MWT<br>baseline | TSUF<br>baseline | NSAA<br>4 y. | 6MWT<br>4 y. | 10MWT<br>4y. | TSUF<br>4y. |
| --- | --- | --- | --- | --- | --- | --- | --- | --- | --- | --- | --- | --- |
| DMD-1 | 4y 0m | M | Quad. | c.6651_6652del; p.Asp221Phefs*3 | 29 | NA | NA | 4.2s | 21 | 443m | 4.5s | 5.6s |
| DMD-2 | 4y 7m | M | Quad. | Duplication Exon 3-7 | 28 | NA | 5.3s | 3.4s | 24 | 473m | 4.8s | 4.7s |
| DMD-3 | 3y 1m | M | Quad. | Deletion Exon 48-55 | 32 | NA | 3.5s | NA | 32 | 515m | 3.2s | 1.9s |
| DMD-4 | 2y 2m | M | Quad. | Deletion Exon 45-50 | NA* | NA* | NA* | NA* | NA | NA | NA | NA |
| DMD-5 | 3y 2m | M | Quad. | c.4918_4919delinsTG | 30 | NA | 5.4s | 3.2s | 31 | NA | 3.4s | 2.9s |
| DMD-6 | 3y 2m | M | Quad. | c.5899C>T; p.Arg1967X | 32 | NA | NA | 4.0s | 34 | NA | 3.5s | 2s |
| DMD-7 | 4y 8m | M | Quad. | Deletion Exon 46-52 | 21 | NA | 5.8s | 3.3s | 29 | 512m | 3.1 | 2.5s |
| Control-1 | 8y | M | Quad. |  |  |  |  |  |  |  |  |  |
| Control-2 | 5y | M | Quad. |  |  |  |  |  |  |  |  |  |
| Control-3 | 9y | M | Quad. |  |  |  |  |  |  |  |  |  |
| Control-4 | 10y | M | Quad. |  |  |  |  |  |  |  |  |  |
| Control-5 | 10y | M | Quad. |  |  |  |  |  |  |  |  |  |

### Characterization of cell populations identified in the skeletal muscle samples

A total of 30857 nuclei from controls and 25817 nuclei from DMD were included in the analysis. Unsupervised clustering using the Seurat package identified 19 different nuclei clusters (Supplemental figure 2)^24^. An analysis of the differentially expressed gene signatures allow us to attribute clusters to 10 putative identities (Figure 1A and Supplemental figure 3). The largest number of nuclei in the samples were from myofibers expressing *Ckm*, a marker of mature myonuclei. As expected, we identified fast and slow type myofibers characterized by the expression of *Myh1* and *Myh2* and, *Myh7b* respectively, but also regenerative fibers characterized by the expression of *Myh3* and *Myh8* (Figure 1 and Supplemental figure 4). Fast myofibers included type IIa fibers expressing *Myh2* and type IIx myofibers expressing *Myh1*, but we just identified a minority of nuclei expressing *Myh4* which is typical of type IIb myofibers. Fast myofibers cluster displayed high levels of genes encoding the fast isoforms of troponins (*Tnnt3*, *Tpm1* and *Tnni2),* the sarcoplasmic reticulum-calcium ATPase 1 (SERCA1/ *Atp2a1*) and glycolytic enzymes (*Eno3, Pfkm, Pkm* and *Pygm)*^25^. The slow myofibers cluster displayed high levels of genes encoding the slow isoforms of troponins (*Tnnt1*, *Tpm3 and Tnni1),* slow myosin light chains *(Myl2* and *Myl3)* and the sarcoplasmic reticulum-calcium ATPase 2 (SERCA2/ *Atp2a2*). Regenerative fibers were identified by the expression of *Myh3* and *Myh8* that encode the embryonic and perinatal MyHC isoforms respectively and, also *Tnnt2* that encodes for an isoform of Troponin-T expressed by cardiac muscle but also embryonic skeletal muscles^26^. As expected, regenerative fibers expressed high levels of *Ncam1*, involved in the reinnervation process of new fibers, and *Myog,* which encodes myogenin and is expressed by both fusing myoblast and new regenerated fibers^27^ . Closely located to the regenerating fibers in the UMAP, we identified a cluster of cells expressing *Pax7* that was identified as satellite cells and shared many genes with regenerative fibers in our samples. This expression profile similitudes illustrated by UMAP is compatible with the origin of regenerative fibers from satellite cells and lead us to study potential gene expression profiles driving cellular transitions using pseudotime trajectories as shown in Supplemental figure 5. As observed cells in the earliest stages of myoblast differentiation showed expression of genes related with ribosomal and mitochondrial function such as *Rpl30*, *Rpl37*, *Mt-co2* or *Mt-co3*, reorganization of the cytoskeleton such as *Myl2*, *Myl1* or *Acta1* and, genes promoting the differentiation of satellite cells such as *Meg3* or *Rassf4*^28, 29^. These were followed by myonuclei expressing genes expressed in fast fibers such as *Myh1*, *Tpm1*, *Tnni2* and, finally, genes expressed in slow fibers such as *Myh7*, *Tnnt1 orTpm3*. Interestingly, we did not identify any population expressing genes related with the myotendinous junction, such as *Col22a1* or *Ankrd1*, as reported in snRNAseq studies done with murine samples that use the whole muscle for the analysis^16, 30, 31^. Moreover, we did not observe a specific cluster expressing genes of the neuromuscular junction (NMJ) as has also been reported in mice, although the expression of *Chrna1* or *Chrng*, was enriched in the regenerative fibers suggesting that there is an active process of remodelling of the NMJ in this cluster as has already been suggested (Supplemental figure 4).^11^

**Figure 1:**
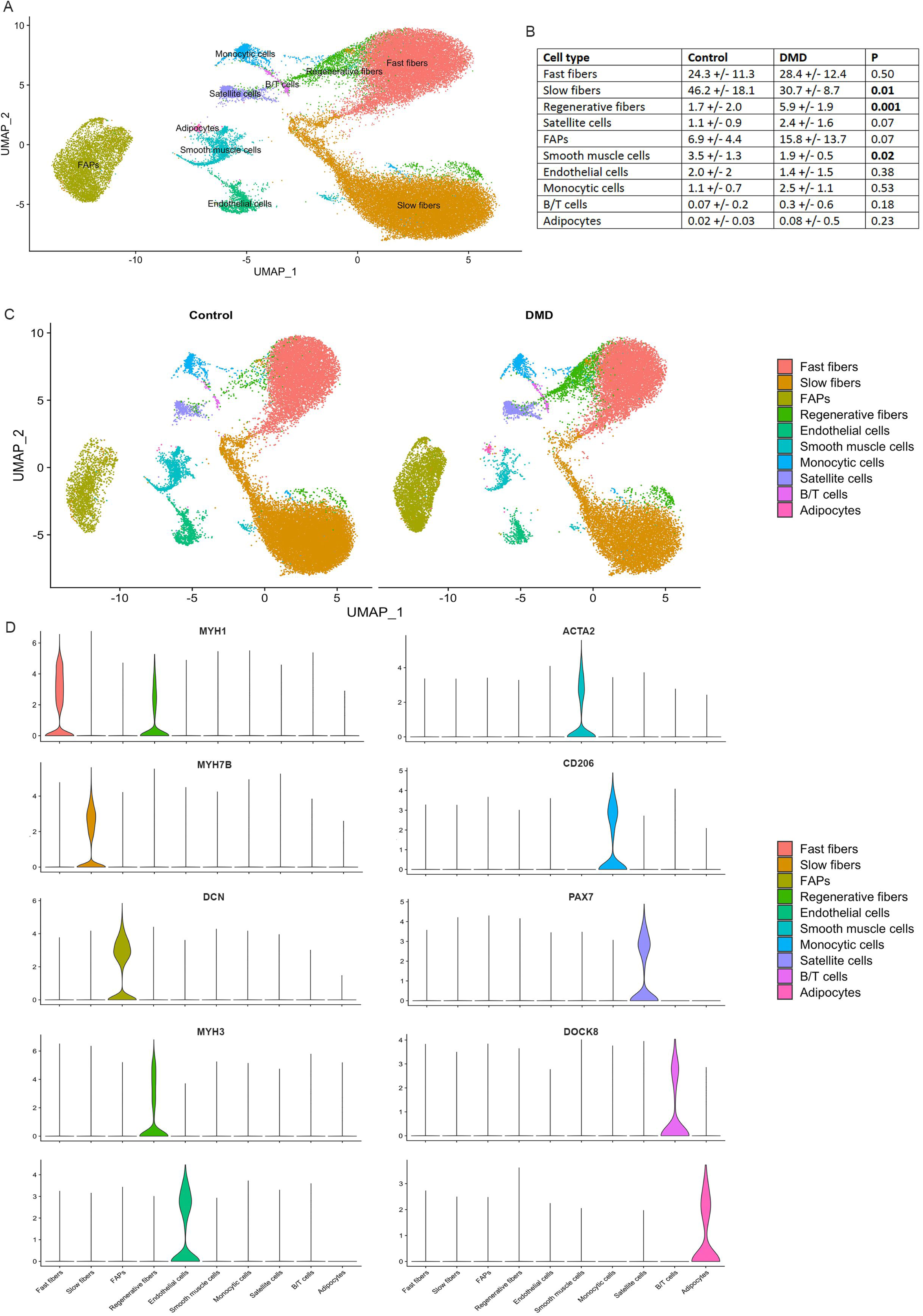
Classification of nuclei/cell types in normal and DMD muscle samples. A) UMAP visualization of all the nuclei from control and DMD samples coloured by cluster identity. B) Table comparing the proportion of cell population between control and DMD samples. C) UMAP showing clusters identified in control (left) and DMD (samples). D) Violin plots showing the expression of selected markers genes for each cluster of nuclei. FAPs: fibroadipogenic progenitor cells.

We observed six other non-myofiber clusters of nuclei that included endothelial cells expressing *Pecam1* and *Ptprb*, smooth muscle cells (SMC) expressing *Acta2*, *Pdgfrb* or *Myh11* or adipocytes expressing *Adipoq* (Figure 1). Inflammatory cells characterized by the expression of *Ptprc/Cd45*, were further divided in macrophages expressing *Mrc1*/*Cd206* and B/T cells expressing *Dok8*^32, 33^. Within the macrophages we identified nuclei expressing M2 markers such as *F13a1, Fcer2/Cd23* and *Cd209* and nuclei expressing M1 markers such as *Cd44*, *Cd86*, *Tlr2*, and *Fcgr3a*^34^. Profibrotic genes, such as *Tgfb1* and *Spp1* were also expressed by M2 macrophages. A large cluster of nuclei was characterized by the expression of *Pdgfra* and *Dcn* which are well-known markers of fibroadipogenic progenitor cells being labelled as fibroadipogenic precursor cells (FAPs) and will be described later in detail.

### Distinctive signature profile in skeletal muscle of patients with DMD

We identified significant differences in the proportion of each cell population when comparing the samples from control and DMD patients (Figure 1B and C). Specifically, there was a significant decrease in the percentage of slow type I fibers (50.9% in controls vs 31.2% in DMD, p=0.01, Mann-Whitney test) and SMC (3.3% vs 1.6%, p=0.02, Mann-Whitney test) and an increase in the percentage of regenerative fibers (1.3 vs 5.7, p=0.001, Mann-Whitney test) in DMD samples. There was also a trend for an increase in the number of satellite cells (0.9% vs 2.4%, p=0.07) and of FAPs (5.6% vs 14.1%, p=0.07). When assessing these differences between samples at the individual level, we observed that the proportion of cells in the muscle was similar among all controls, while there was greater variability in the DMD samples, which is compatible with an active process of muscle degeneration going through different stages (Figure 2A and B). PCA analysis differentiated between DMD and controls by assessing the proportion of each cell population in the biopsy (Figure 2C). The control patients were all closely located in the 2D dimension PCA map. Interestingly, the distribution of DMD patients on the PCA graph moved according to their clinical phenotype revealing two different subgroups, one that could be earlier in the process of muscle degeneration and included five samples from younger patients (2 to 3 years) with a mildly affected muscle function and characterized by an increase in the number of nuclei from regenerative fibers (DMD-3 to DMD-7) and, another group of two samples from patients slightly older (4 years) who were already showing clear signs of muscles weakness and were more advanced in the process of muscle degeneration with a reduced number nuclei from slow and fast fibers and an increase in the nuclei from FAPs (DMD-1 and DMD-2).

**Figure 2:**
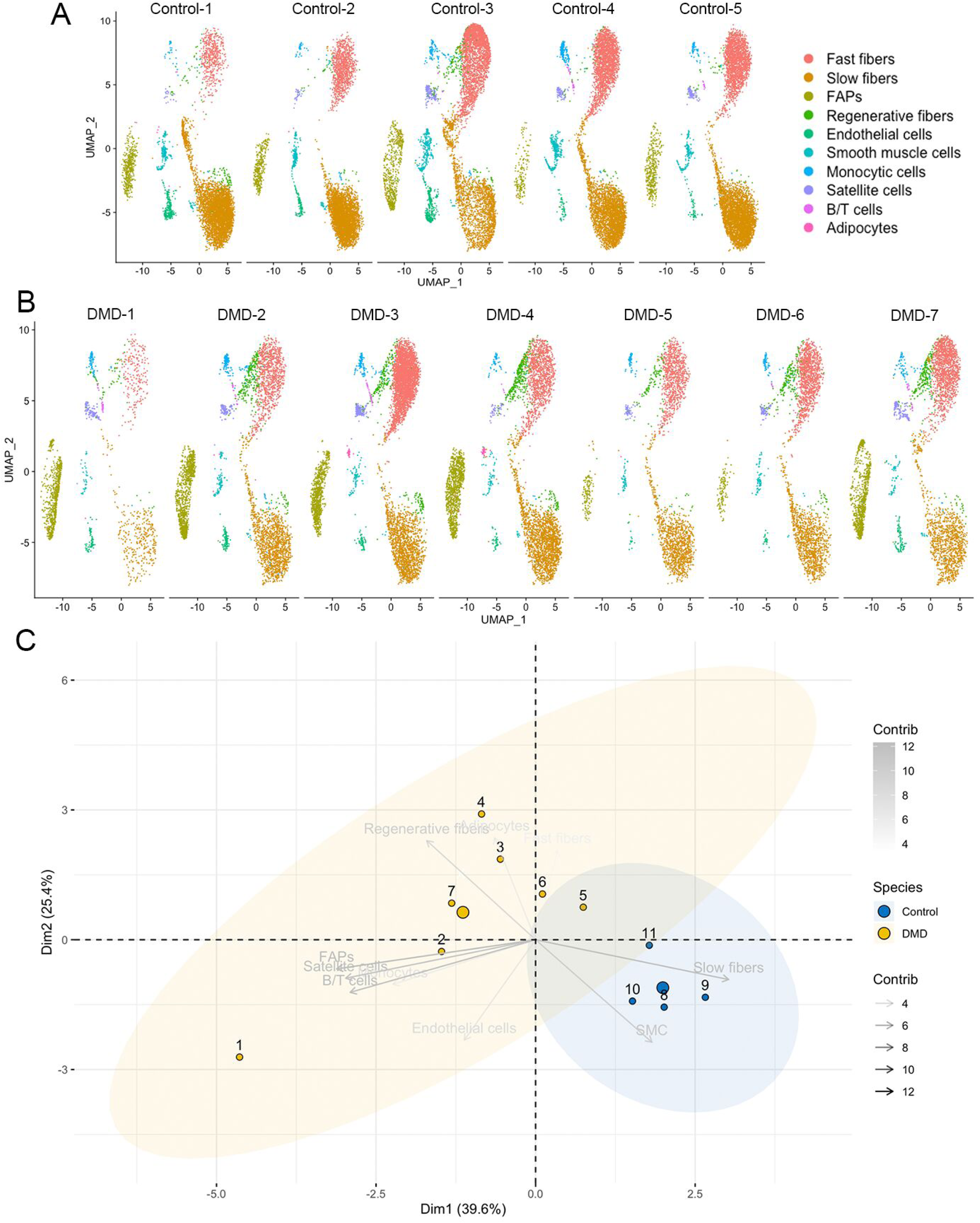
Differences in cell population to the individual level. A) UMAP visualization of nuclei from control individuals coloured by cluster identity. B) UMAP visualization of nuclei from DMD individuals coloured by cluster identity. C) PCA analysis showing the distribution of individuals based on each cell population proportion.

### Differential expression of genes in muscle fibers of patients with DMD

To gain insight in myofibers transcriptome in each cluster, we compared the gene expression profile of fast and slow muscle fibers nuclei between healthy controls and DMD patients (Figure 3). Considering those genes with a higher log_2_FC>0.5, we found 292 genes significantly upregulated and 85 downregulated in fast myofibers while in slow fibers we found 238 upregulated and 89 downregulated in DMD compared to controls. Fast and slow myofibers in DMD shared several genes in the top ten upregulated genes, such as *Myh3*, a characteristic marker of regenerative myofibers (log_2_FC=3.1 and 2.1 respectively), *Meg3*, a LncRNA involved in myoblast plasticity and differentiation (log_2_FC=2.3 and 2.9 respectively) and *Ldb3* which acts as an adapter in skeletal muscle to couple protein kinase C-mediated signaling via its LIM domains to the cytoskeleton (log_2_FC=1.6 and 1.3 respectively) (Figure 3 A-D)^35, 36^. Interestingly, we observed an upregulation of genes involved in the transport of calcium (*Cacnas1*, *Ryr1*) and also, proteases such as *Capn3 and Capn2*, two pathways already known to be involved in the process of muscle degeneration in DMD^37^. Gene set enrichment analysis (GSEA) revealed several dysregulated molecular pathways (Figure 3H). We observed an enrichment in both slow and fast fibers of the expression of genes involved in myogenesis, and muscle growth compatible with an active process of muscle regeneration, but also genes involved in axon guidance suggesting that new and regenerated muscle fibers release signals for the growth of terminal axons needed for reinnervation or, genes involved in adherens junction probably due to the need of new fibers to link again to the ECM. Interestingly, DMD fibers had a reduction in several metabolic pathways, when compared with control individuals especially oxidative phosphorylation but also glycolysis and lipid transport confirming previous observations^11^.

**Figure 3:**
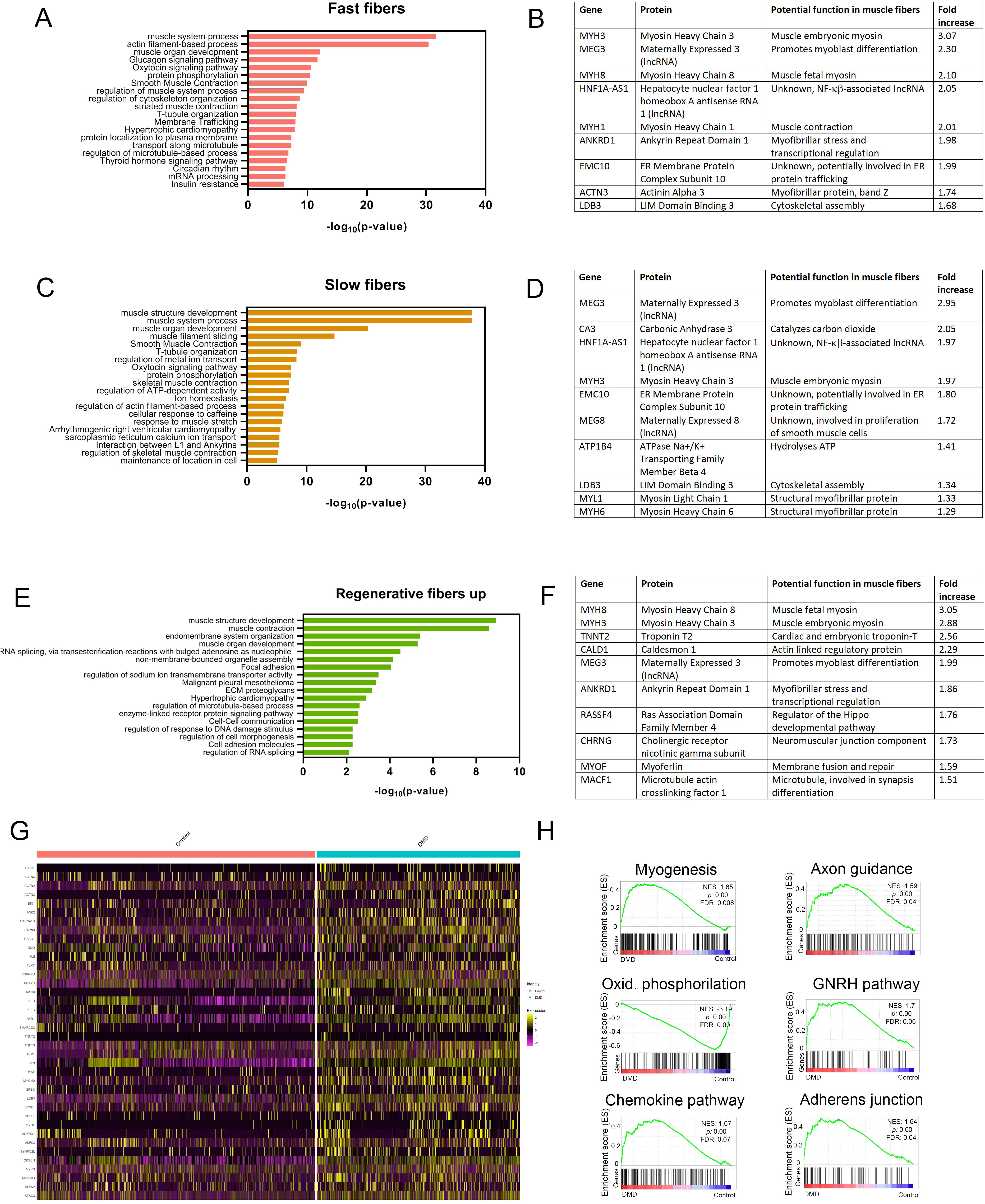
Analysis of gene expression changes in DMD myonuclei compared to control myonuclei. A) Top molecular pathways upregulated in fast fibers. B) List of the top ten genes upregulated in myonuclei of fast fibers of DMD samples. C) Top molecular pathways upregulated in slow fibers. D) List of the top ten genes upregulated in myonuclei of slow fibers of DMD samples. E) Top molecular pathways upregulated in regenerative myofibers. F) List of the top ten genes upregulated in myonuclei of regenerative myofibers. G) Heatmap showing expression of genes involved in muscle growth in Control and DMD fast and slow myonuclei. H) GSEA plots showing enrichment score (ES) of the significant enriched hallmark gene sets in fast and slow myonuclei. A positive value of ES indicated enriched in DMD and a negative value indicates enriched in normal condition and down-regulated in DMD. GSEA: gene set enrichment analysis; NES: normalized enrichment score; FDR: false discovery rate. Oxid: Oxidative. GNRH: Gonadotropin hormone-releasing hormone

In the case of regenerative myofibers, as they were just a minority in control patients (1.7%) and much more abundant in DMD (5.9%), we decided to analyze the genes increased in this population compared to slow and fast muscle fibers of patients with DMD. We found 69 upregulated and 63 downregulated genes in regenerative fibers. As expected, among the top ten upregulated genes we identified many involved in the process of muscle regeneration such as *Myh3*, *Myh8*, *Tnnt2* or *Cald1* involved in the regeneration of the myofibrillar system, but also genes involved in membrane fusion (*Myof*), neuromuscular junction development (*Chrng* or *Macf1*) or Hippo pathway (*Rassf4*) involved in myoblast activation and muscle growth^38, 39^.

As myogenesis and cell growth was one of the main pathways upregulated in muscle fibers, we further investigated the mechanism controlling these processes^40^. We analyzed several pathways and as shown in supplemental figure 6, we observed an upregulation of the MEF2 family of transcription factors that have been classically involved in the myogenic program, especially *Mef2a* and *Mef2c*, while *Myf6* expression, an inhibitor of MEF2 was reduced in the muscle fibers^41^. Additionally, expression of *Tead1* and *Tead4*, the downstream effectors of the Hippo pathway involved in muscle hypertrophy, were also upregulated in DMD muscle fibers, associated with an increase in *Yap* and *Taz* expression two cofactors of this pathway^42^. In concordance with these findings, pro-atrophic genes such as *Murf1 (Trim63)* and *Atrogin-1* were downregulated in slow/fast muscle fibers. We did not observe an increase in the expression of genes belonging to the myostatin or insulin growth factor pathway in slow or fast myofibers.

### FAPs from patients with DMD express genes related with cell proliferation and extracellular matrix remodelling

We compared the transcriptional profile of FAPs from healthy controls and DMD patients. DMD FAPs had a significant upregulation of 249 genes and a downregulation of 68 genes (log_2_FC>0.5) compared to controls. Among the top upregulated genes, we found genes encoding different type of collagens (*Col1a1, Col1a2, Col6a6, Col3a1* or *Col21a1* among others), but also other components of the extracellular matrix such as elastin (*Eln*) and several fibulins. Genes encoding for proteins involved in extracellular matrix assembly such as *Sned1*, matrix remodeling (*Adamtsl1*) or interaction between cells and matrix such as laminins (*Lamb1* and *Lama4*) were also upregulated^43^. As many of the genes produced by DMD FAPs were components of the extracellular matrix, we compared the expression levels of matrix components between control and DMD samples and observed significant differences, not only in the expression levels but also in the components identified as shown in supplemental figure 7. Apart of the extracellular matrix genes, we observed an upregulation of genes involved in relevant signaling pathways such as PDGF and NCAM signaling, tyrosine kinase activation and, Rho-GTPase cycle suggesting that FAPs are not a mere producer of extracellular matrix but they could also play a role as a potential regulator of the activity of other muscle resident cells (Figure 4B and D).

**Figure 4:**
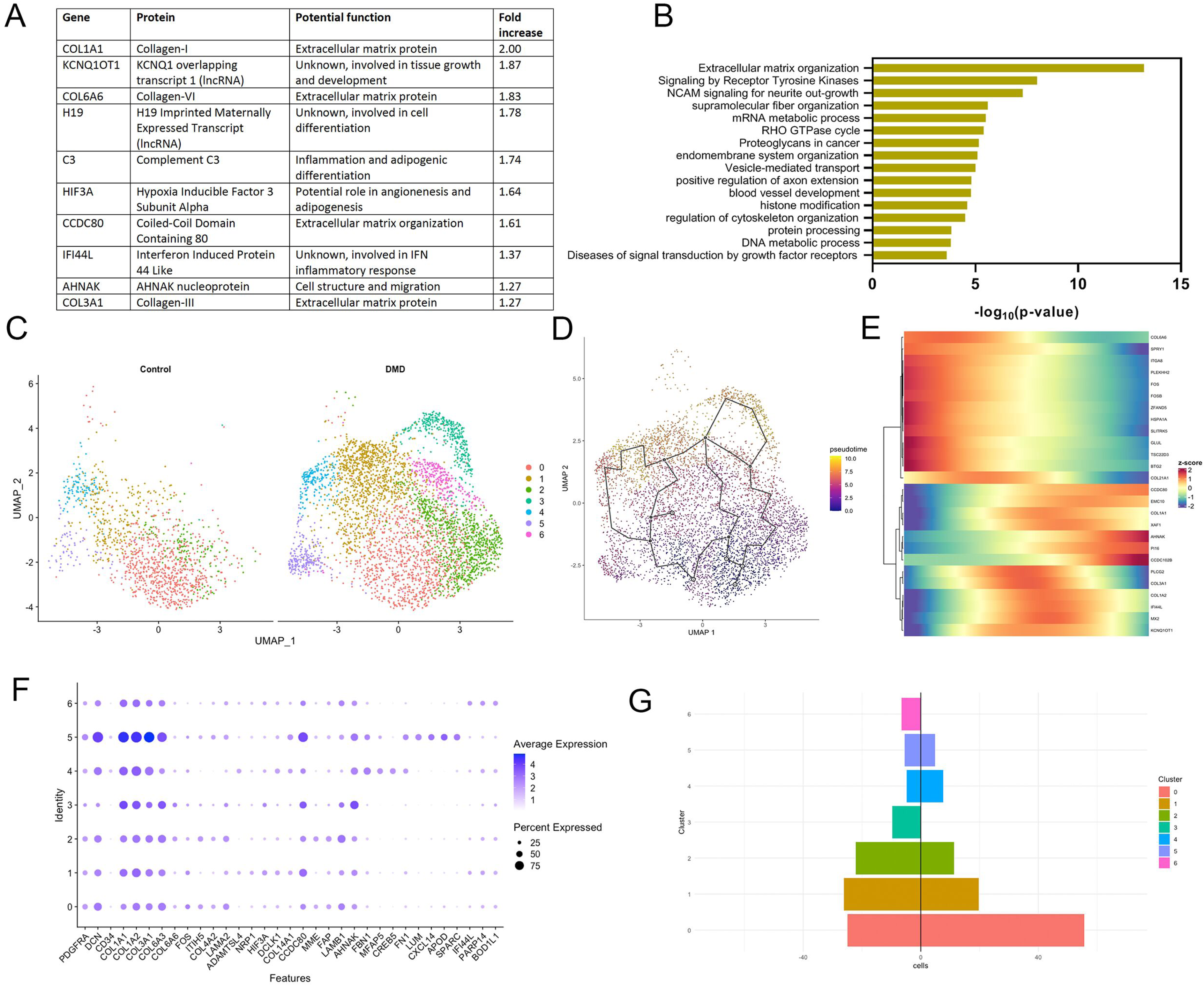
Analysis of gene expression changes in DMD FAPs compared to control nuclei. A) List of the top ten genes upregulated in FAPs of DMD samples. B) Top molecular pathways upregulated in DMD FAPs. C) UMAP visualization of nuclei from FAPs of control and DMD individuals coloured by subpopulation identity. D) Monocle analysis showing pseudotime trajectories of the re-clustered FAPs. E) Heatmap showing selected gene expression across pseudotime trajectories. F) Selected genes expressed in each FAP subcluster. G) Population of subclusters of FAPs in Control and DMD samples.

**Figure 5:**
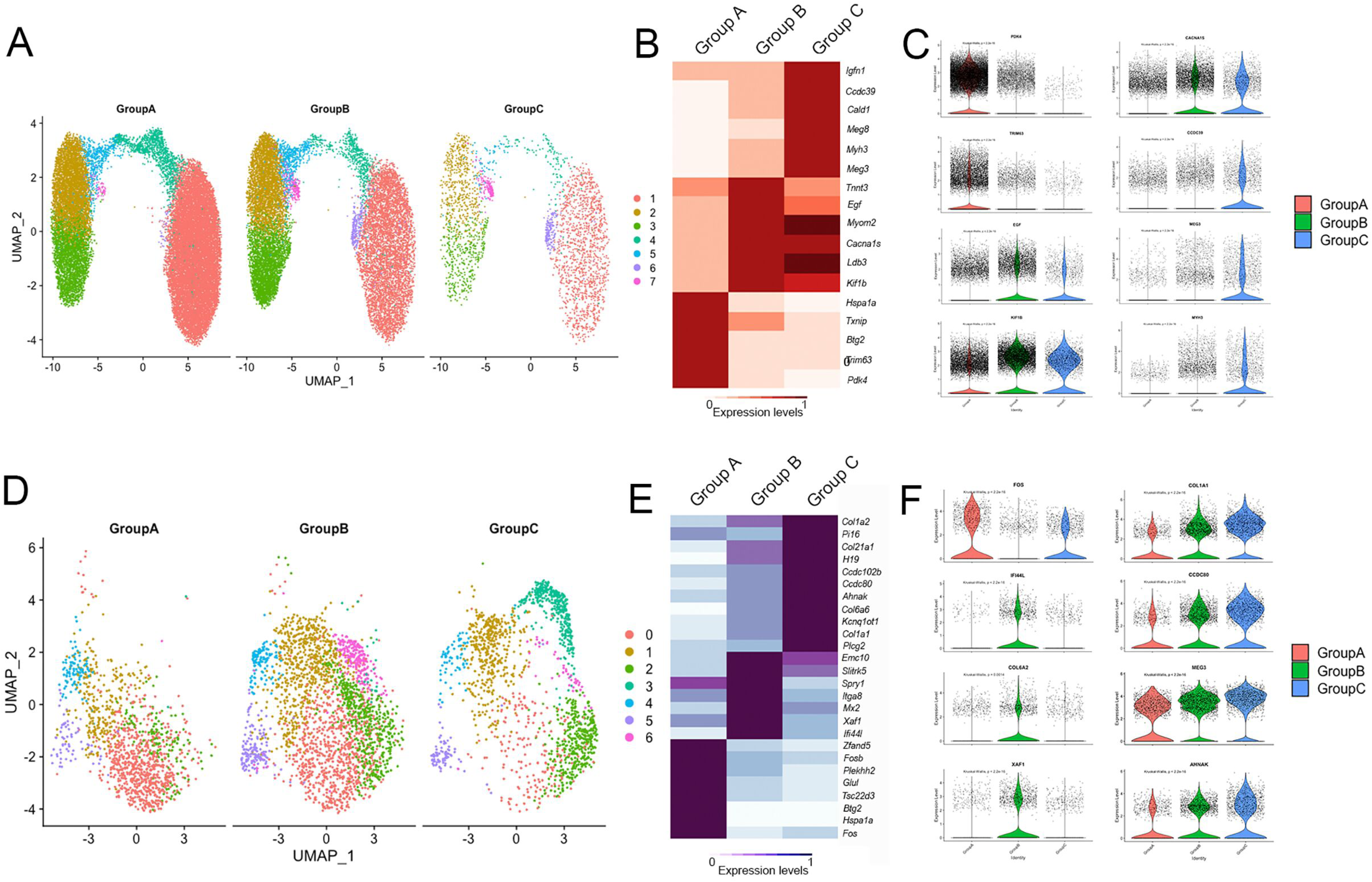
Differences in cell population and gene expression profile between stable and progressing DMD patients. A) UMAP of the subpopulations of myonuclei in controls, stable and declining DMD patients. B) Heatmap showing the top upregulated genes in myonuclei from controls, stable and declining DMD patients samples. C) Violin plot showing the expression of selected markers genes for myonuclei of controls, stable and declining DMD patients. C) UMAP of the subpopulations of FAPs in controls, stable and declining DMD patients. B) Heatmap showing the top upregulated genes in FAPs from controls, stable and declining DMD patients samples. C) Violin plot showing the expression of selected markers genes for FAPs of controls, stable and declining DMD patients.

Based in the myriad of biological processes upregulated in DMD samples, we decided to explore if there could be subpopulations of FAPs at different states of differentiation that could be distinguished based on their gene expression profile. We re-clustered the FAP subgroup and identified seven different subclusters of cells as shown in Figure 4C. These clusters shared the expression of many genes such as *Dcn*, *Pdgfra*, *Col1a1* or *Col3a1*, however some genes that were preferentially expressed in some of the clusters (Figure 4F). For example, cluster 0 that was predominant in control samples expressed higher levels of the antiproliferative genes *Fos*(*p55*) and *Itih5*, but also *Col4a2* and *Lama2*. Cluster 4 was characterized by the expression of *Fbn1* while cluster 5 was characterized by the expression of *Lum* and could confirm the existence of this population of FAPs in DMD patients recently described by Rubinstein et al^44^. Lum+ FAPs were the ones expressing the highest levels of collagen related genes such as *Col1a1* or *Col3a1*. Two of the clusters identified, cluster 3 and 6, were almost exclusively present in DMD samples and were characterized by the expression of genes involved in cell proliferation such as *Ahnak*, *Ccdc102b* and *Podn* or *Parp14*, *Bod1l1* or *Smg1* respectively. Interestingly, cluster 6 was distinctively present in the patient that had the greatest decline in muscle function during follow up. Monocle analysis identified potential trajectories in the differentiation process of FAPs over time that started in Cluster 0, majoritary of controls, and end in Cluster 6 (Figure 4D). Moreover we also identified genes differently expressed through the differentiation process (Figure 4E).

As expected, the population of adipocytes, a type of cell known to derive from FAPs in the skeletal muscles, was higher in DMD samples than in controls. Further, we studied the molecular pathways activated in adipocytes based on their gene expression profile and observed that apart of pathways involved in lipid metabolism, adipocytes had an increased expression of genes of the Rho pathway and genes encoding for components of the basal lamina such as *Col4a1*, *Col4a2*, *Lama4* and, *Lamb1* (Supplemental figure 8)

### Gene expression in different stages of disease progression

We were interested in investigating differences in the gene expression profile between DMD patients at different stages of disease progression. To do so, we reviewed the clinical and muscle function information present in the clinical notes and observed that there were consistent differences in clinical function at baseline and disease progression over the first four years after the muscle biopsy was obtained. As show in table 1, patients DMD 3, 5, 6 and 7 had mild muscle impairment at baseline and showed either stabilization or improvement in muscle function during follow-up period. On the other hand, patients DMD 1 and 2, showed a worse baseline performance and a decline in muscle function during the follow up period^45^. We explored if there were differences between control samples (Group A), stable patients (Group B) and declining patients (Group C) in the gene expression profile of muscle fibers and FAPs. Muscle fibers from controls were enriched in the expression of genes such as *Pdk4 and Txnip* involved in the metabolism of glucose and lipids, *Linc-Pint* and *Btg2* inhibiting cell division and, *Trim63* (*Murf1*) involved in protein ubiquitination^46-48^. In the case of DMD patients, we did not observe many differences between Group B and C in the upregulated genes of, that were predominantly involved in muscle regeneration, either on satellite cell activation, membrane fusion or sarcomere assembly (*Meg3*, *Meg8*, *Myh3, Cald1, Igfn1, Myof or Myo18B*). However, when we analyzed gene expression of FAPs among the three groups we did observe interesting results. As previously mentioned, control FAPs had a statically significant upregulation in the expression of antimitotic genes, such as *Fos* or *Btg2* compared to DMD FAPS. FAPs from Group B (DMD stable patients) had a statically significant upregulated expression of the proapoptotic gene *Xaf1*, interferon induced genes such as *Ifi44l* or *Mx2* or the profibrotic differentiation transcription factor *Spry1* while, FAPs from Group C (DMD declining patients) had the highest expression of collagen genes (*Col1a1*, *Col1a2*, *Col3a1*) but also high expression of genes actively involved cell division (*Ccdc80* or *Ccdc102b*), indicating that in the declining patients FAPs actively proliferate and express EXM components replacing the muscle fibers lost.

### Communication between cells populations is dysregulated in DMD muscles

We studied the predicted intercellular communications of each cell population and compared the communication network between control and DMD using CellChat package^49, 50^. The analysis revealed significant differences in the number of cell interactions. As shown in Figure6A and B, FAPs and satellite cells became the most important source of ligands in DMD, potentially interacting with all other cell populations. Adipocytes, which were mainly present in DMD samples, played also an important role in cell to cell communication in DMD samples. CellChat detected 55 significant ligand-receptor pairs in the Control dataset and 61 in the DMD samples among the 11 nuclei clusters (Figure 6 C-D). A number of molecular pathways were identified exclusively in DMD samples such as cadherins (CDH), NCAM, major histocompatibility class-I and neuroregulin, while other were exclusively identified in Control samples including CD40, CD80 or IL-2 among others. Signalling pathways upregulated in DMD samples were involved in several processes such as cell migration and remodelling of extracellular matrix (FGF, Collagen, Laminin), nerve growth and reinnervation (NCAM, NGF and NPR2) and inflammation (MHC-I, CXCL, THBS). As FAPs were identified as the main producer of ligands outgoing to other cell populations both in control and DMD (Figure 6E and F) we decided to investigate further the main molecular signals released by these cells (Figure 6G).

**Figure 6:**
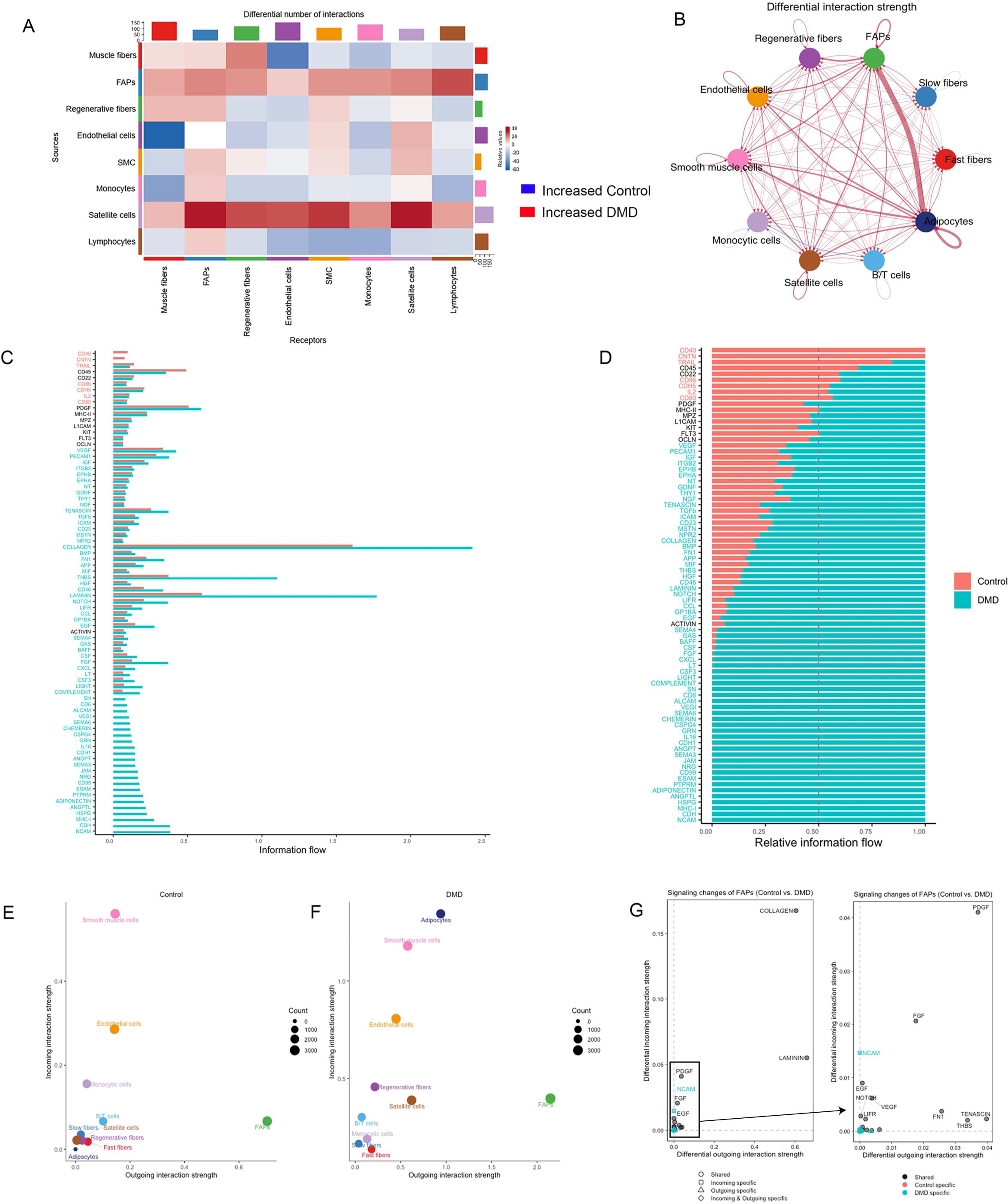
Analysis of intercellular communications in control and DMD muscle samples. A) Heatmap showing differential number of interactions between clusters in DMD samples compared to controls. Red: increased interactions in DMD. Blue: Increased interactions in controls. B) Chord plot displaying intercellular ligand-receptor interaction strength comparing DMD and control samples. Red: increased interactions in DMD. Blue: Increased interactions in controls. C) Bar graph showing relative information flow per each signalling path in control(red) and DMD (green) samples. D). Bar graph showing weight of each signalling path in control (red) and DMD (green) samples. E) Dot plot showing the weight of each cell cluster in outgoing-incoming signalling in control samples. F) Dot plot showing the weight of each cell cluster in outgoing-incoming signalling in control samples. G) Dot plot showing the main molecules released and received by FAPs in DMD and control samples.

Collagens and laminins were the most upregulated molecules signals produced by FAPs in DMD, followed by others such as members of the PDGF and FGF family, but also tenascin, thrombospondin and fibronectin. As collagen and laminin were components of the ECM and could potentially influence all muscle cell behaviour, we decided to study more precisely their potential communication network. Network centrality analysis of the inferred collagen identified FAPs as the most prominent sources of collagen either in control and in DMD samples, acting onto endothelial and smooth muscle cells on control, but also onto adipocytes in DMD (Figure 7). Notably, among all known ligand-receptor pairs, DMD collagen signalling was mainly dominated by collagen I, IV and VI and its receptor *Itga1/Itga2* + *Itgb1*. FAPs were also the most prominent source of laminin either in control and DMD samples, although adipocytes became an important source as well in DMD (Figure 8). In control samples, laminin pathway was dominated by the *Lama2* and *Lamb1* ligands and its *Itga1/Itgb1* and *Itga7/Itgb1* receptors on endothelial and smooth muscles cells. In DMD, *Lama4* and *Lamb1* predominated acting through multiple *Itga/Itgb* receptors on adipocytes, regenerative fibers and adipocytes in addition to smooth and endothelial cells.

**Figure 7:**
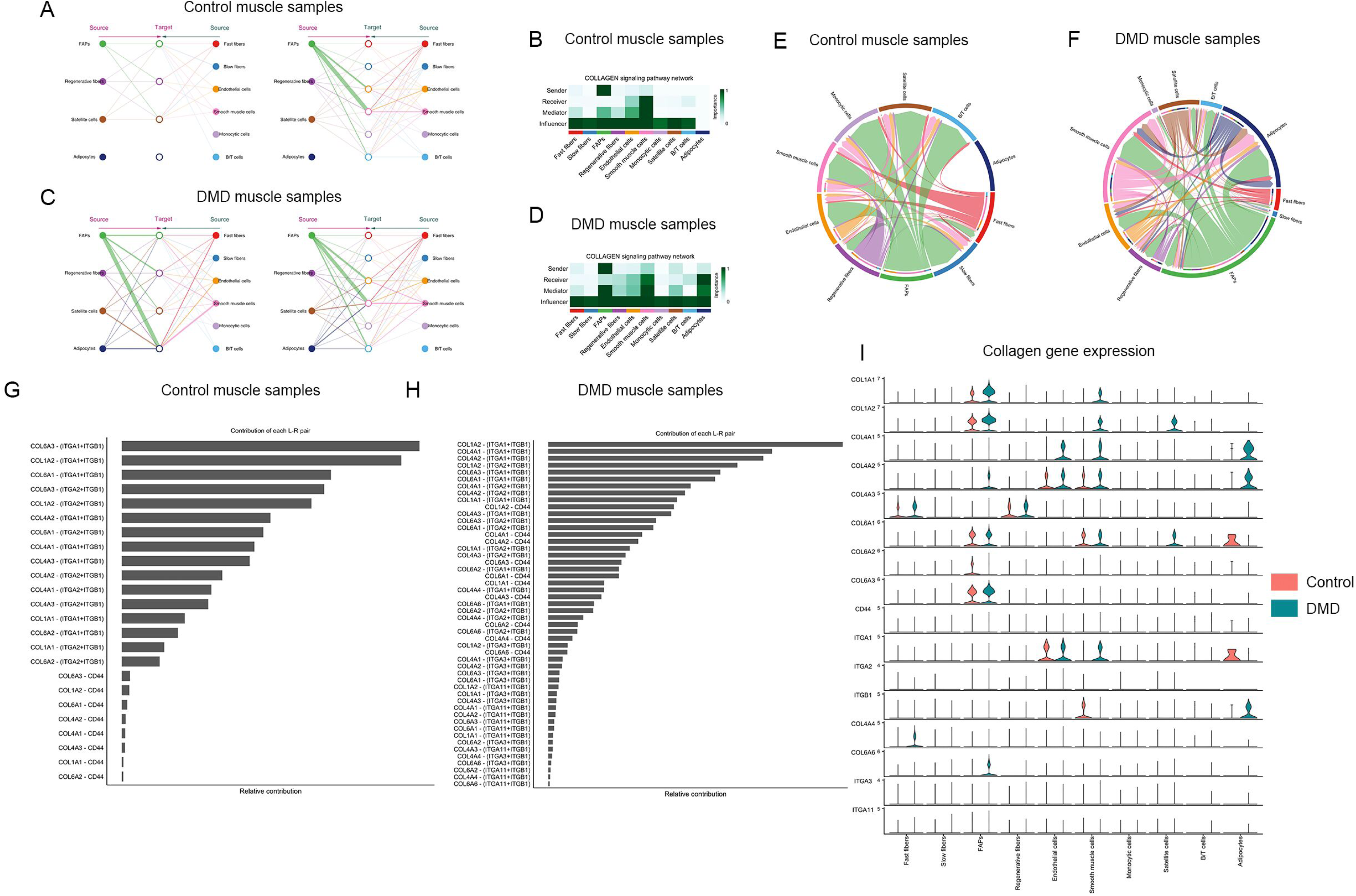
Collagen signalling pathway in control and DMD muscle samples. A) Hierarchical plot shows the inferred intercellular communication network for collagen signalling. This plot consists of two parts: Left and right portions highlight the autocrine and paracrine signalling to FAPs, regenerative fibers, satellite cells and adipocytes and to the other clusters identified, respectively. Solid and open circles represent source and target, respectively. Circle sizes are proportional to the number of cells in each cell group and edge width represents the communication probability. Edge colours are consistent with the signalling source. B) Heatmap shows the relative importance of each cell group based on the computed network centrality measures of collagen signalling network in control samples. C) Hierarchical plot shows the inferred intercellular communication network for collagen signalling in DMD samples. D) Heatmap shows the relative importance of each cell group based on the computed network centrality measures of collagen signalling network in control samples. E) Chord plot displaying intercellular communication network for collagen signalling in controls. F) Chord plot displaying intercellular communication network for collagen signalling in DMD. G) Relative contribution of each ligand-receptor pair to the overall communication network of collagen signalling pathway in control samples, which is the ratio of the total communication probability of the inferred network of each ligand-receptor pair to that of collagen signalling pathway. H) Relative contribution of each ligand-receptor pair to the overall communication network of collagen signalling pathway in DMD samples. I) Violin plot showing the expression distribution of signalling genes involved in the inferred collagen signalling.

**Figure 8:**
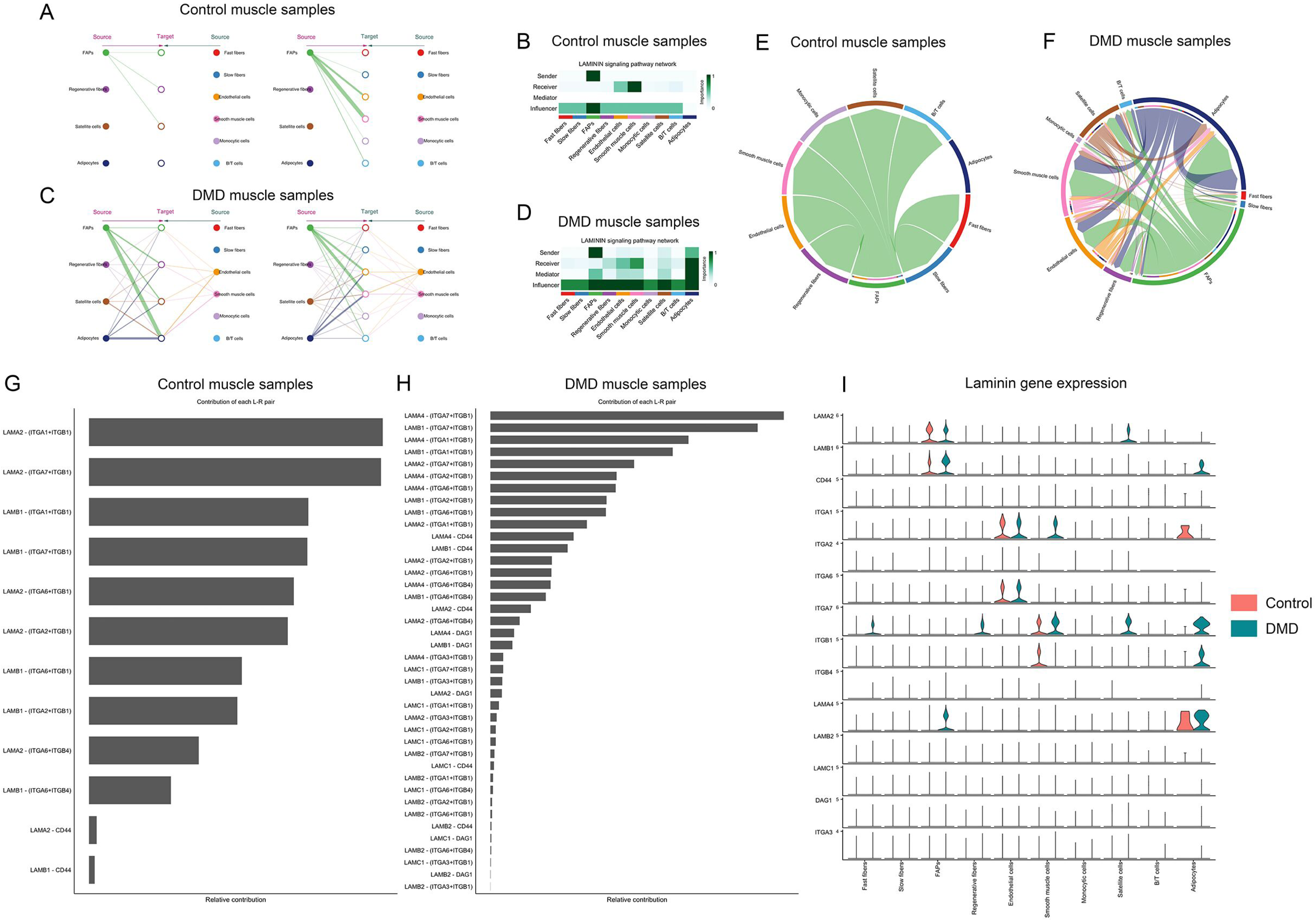
Laminin signalling pathway in control and DMD muscle samples. Hierarchical plot shows the inferred intercellular communication network for laminin signalling. This plot consists of two parts: Left and right portions highlight the autocrine and paracrine signalling to FAPs, regenerative fibers, satellite cells and adipocytes and to the other clusters identified, respectively. Solid and open circles represent source and target, respectively. Circle sizes are proportional to the number of cells in each cell group and edge width represents the communication probability. Edge colours are consistent with the signalling source. B) Heatmap shows the relative importance of each cell group based on the computed network centrality measures of laminin signalling network in control samples. C) Hierarchical plot shows the inferred intercellular communication network for laminin signalling in DMD samples. D) Heatmap shows the relative importance of each cell group based on the computed network centrality measures of laminin signalling network in control samples. E) Chord plot displaying intercellular communication network for laminin signalling in controls. F) Chord plot displaying intercellular communication network for laminin signalling in DMD. G) Relative contribution of each ligand-receptor pair to the overall communication network of laminin signalling pathway in control samples, which is the ratio of the total communication probability of the inferred network of each ligand-receptor pair to that of laminin pathway. H) Relative contribution of each ligand-receptor pair to the overall communication network of laminin signalling pathway in DMD samples. I) Violin plot showing the expression distribution of signalling genes involved in the inferred laminin signalling.

## 3. Discussion

We have investigated the gene expression of cells present in muscle samples of DMD patients and age/gender matched controls. Our study has revealed novel and significant differences in the population of cells present in DMD muscles, their gene expression profile identifying several dysregulated molecular pathways and changes in the intercellular signalling network. Our data show that these cellular and genetic modifications are dynamic through disease natural history impacting patient’s muscle function and clinical progression.

One of the most relevant findings is the change on the proportion of each cell population in the muscle samples of DMD patients happening from early stages of the disease. The most consistent change is the increase in the number of regenerative fibers that probably contribute to maintain muscle function when there is not yet a massive loss of mature fibers, as observed in our patients who had mild muscle function impairment and who were clinically stable over four years after the biopsy was obtained. However, despite a regenerative gene signature was also observed in the muscle samples of the declining patients, they had reduced number of mature muscle fibers, an increase in the number of FAPs and presence of adipocytes. These data linking proportion of cell types and muscle function, suggest that the cellular changes observed in skeletal muscles of patients with DMD are identified since early ages and are dynamic over time influencing disease progression^51, 52^. Further studies analysing muscle samples from patients at later stages of disease progression are required to better understand the complexity of how the cell population changes over time and which molecular pathways are progressively activated. However, the availability of muscle biopsies at these later stages of disease progression is usually very restricted or absent because once the diagnosis is reached, muscle biopsies are not requested.

In concordance with an increased number of nuclei corresponding to regenerative fibers, we observed an upregulation of genes involved in myogenesis and muscle repair in myonuclei of slow and fast muscle fibers suggesting that at these early stages of the disease, injured muscle fibers are efficiently activating the regenerative machinery expressing genes coding for several developmental isoforms of sarcomeric proteins, molecules involved in linking new fibers to the extracellular matrix or molecules involved in the reorganization of the T-tubule system. Consequently, genes involved in muscle fiber growth and hypertrophy were upregulated in slow/fast myonuclei, while proatrophic factors were downregulated^53^. Interestingly, genes codifying components of the neuromuscular junction were also upregulated, including the fetal nAChR gamma subunit (*Chrng*) which expression is stopped at birth and substituted by postnatal epsilon subunit (*Chrne*), reinforcing the idea that neuromuscular junction is remodelled during muscle regeneration^54^. Moreover, genes involved in axon guidance such as *Ncam1*, *Gdnf* and members of the semaforin family (*Sema3a* or *Sema4d*) were also increased suggesting that reinnervation is partially driven by signals released from the newly formed myofibers^53^. Although the regenerative signature was predominant in the myofibers, we also identified upregulation of genes involved in the process of protein degradation, such as overexpression of the proteases *Capn3* and *Capn2* and calcium channels *Ryr1* and *Cacnas1s*, two pathways suggested to be involved in the process of muscle degeneration in DMD^55, 56^. However, and in concordance with a cell that is actively growing, genes coding for enzymes involved in protein ubiquitination or atrophy were repressed, even though they have been reported to be increased in muscle fibers of the mdx mice^21^. These results agree with previously published data using bulk RNA analysis or proteomics of muscle biopsies of DMD patients at early stages of diseases progression, showing a strong muscle regeneration signature and validating the results of our snRNAseq analysis^11, 12^.

Increase in the number of FAPS was another prominent change observed in DMD patients. DMD-FAPs were characterized by an upregulation of the genes involved in expansion and remodelling of the ECM including but not limited to collagen and laminin genes (*Col1a1*, *col6a6* and *col3a1*), metalloproteinases (*Adamtsl1*) and molecules involved in the assembly of the extracellular matrix (*Sned1, Pcolce, Dcn*). Interestingly, we observed significant differences in the expression of genes codifying ECM components between DMD and control samples, suggesting that there are differences not only in the quantity of some of the components but also in the composition of the matrix with a substantial increase in collagen I, III and VI which are part of the interstitial matrix, while collagen IV, one of the components of the basal lamina remained stable^57, 58^. A complete understating of the impact that these changes have in muscle cells’ behaviour is lacking, but it is know that ECM apart of providing structural support to cells, also facilitates communication, regulates cell growth, promotes or restrict cell movement and transmits mechanical signals ^59^. We have identified different subpopulation of FAPs present in both control and DMD muscles, including the already described Lum+ and Fbn1+ cells, but not other FAPs population described in murine models, such as the DPP4+ FAPs^44, 60^. Our analysis revealed a change in the predominant FAP subpopulations present in DMD and the existence of subpopulations that are not present in controls characterized by upregulation of genes involved in cell division. These subpopulations were mainly identified in patients with a declining muscle function during disease progression suggesting that FAPs cell expansion could be a hallmark of muscle degeneration in DMD. However, a complete understanding of the potential role of these subpopulations of FAPs requires further characterization of the cells. Interestingly, FAPs were identified as the main messenger of signals either in control and DMD muscles by CellChat suggesting that FAPs are not a simple producer of ECM. The paracrine signals generated by FAPs targeted mostly endothelial and smooth muscle cells in control samples but also satellite cells, regenerative fibers and adipocytes in DMD. The interaction between these cells is driven by many different molecules, but we have observed that collagens and laminins could play an important role in intercellular communication driven by FAPs in the muscle. Adipocytes, which derive from muscle resident FAPS, irrupts in DMD muscle as an important player regulating cell signalling through the production of laminins contributing to the modified ECM. We have identified a substantial number of signalling pathways predominant or even only observed in DMD samples compared to control, somehow drawing a kind of DMD signalling fingerprint which helps to summarize the events that are taking place in these early stages of muscle degeneration. These events include ECM remodelling but also cell adhesion, migration, chemotaxis of cells, proliferation, differentiation and inflammation.

The clinical features of the patients included in this study was homogeneous, as expected for patients at two, three or four years of age when the muscle biopsy was obtained. However, muscle function revealed subtle differences between patients and two groups were distinguishable, one characterized by mild impaired muscle performance at baseline and stability over time and another one with worse muscle function and deterioration after muscle biopsy. It is important to remark that the muscle biopsied was the quadriceps which is essential for the muscle function tests performed. When we compared differences in the gene expression profile between stable and progressive patients we did not observe major significant differences between myofibers that were characterized by an strong regenerative signature. However, we did observe that FAPs of the declining group were characterized by expression of genes involved in cell division and have an upregulation of genes encoding components of the ECM, compared to stable patients. This suggests that the existence of a population of active proliferative FAPs that could produce higher levels of collagen is key in the process of active muscle degeneration reinforcing the idea that treating patients with drugs inhibiting FAP proliferation or activation could be beneficial for DMD patients^61, 62^.

Up to date our understanding of the process of muscle degeneration in muscular dystrophies is mainly based in studies performed in murine models of the disease^63, 64^ . These studies have provided valuable knowledge, although the results obtained have not been always validated in humans, mainly because of the lack of good animal models mimicking the process of muscle degeneration observed in patients. This is especially true in the case of DMD, where the existing murine models develop a milder disease characterized by loss of muscle fibers and its replacement by fibrotic tissue only at late stages of the disease, while there is almost no fat present in the muscles^65, 66^. The study of disease mechanisms in DMD using human samples can be complex because of the lack of available tissue especially since the popularization of molecular studies for diagnosis purposes^67^.

In summary, we have studied the gene expression profile to the single nuclei level in muscle samples of controls and DMD patients at an early stage of disease progression. We have focused our analysis on changes happening in muscle fibers and FAPs, as they the two population of cells showing more changes in their number between controls and DMD. We have observed a substantial number of genes which expression is dysregulated in both types of cells pointing towards an enhanced regenerative activity in DMD patients at this stage, associated with an increase proliferative activity of FAPS, which produce high levels of extracellular matrix components.

## 4. Material and methods

### Muscle specimens from healthy controls and patients with DMD

Muscle samples of boys with genetically confirmed DMD were obtained for diagnosis purposes or for research from patients seen at the Newcastle Hospital NHS Foundation Trust or at the Hospital Sant Joan de Deu Hospital in Barcelona. Muscle samples from controls were obtained from healthy children undergoing a orthopaedic surgery at Hospital Sant Joan de Deu in Barcelona. Patients’ and controls’ parents or legal representatives signed a consent form for the biopsy. Muscle samples were stored at the biobanks of each institution in liquid nitrogen tanks. The obtention of the biopsy and storage in the biobank was approved by local Ethics Committee at both Institutions. The research study performed here with the samples was approved by the Ethics Committee of the Newcastle University (reference 13866/2020). These samples were studied with conventional staining protocol to confirm that they were normal samples.

### Nuclei purification from human muscle biopsies

Frozen muscle biopsies (25 to 40 mg) were placed in homogenization buffer (0.25M sucrose and 1%BSA in Mg2+/Ca2+-free, RNase-free PBS). Tissue was homogenized using a Tissue Ruptor II (Qiagen) and incubated for 5 min with 2.5% Triton-X100 (added at 1:6 ratio). The resulting homogenates were filtered through 100 μm and 70 μm strainers (Miltenyi Biotec), pelleted by centrifugation (3000 xg, 10 min at 4°C), resuspended in sorting buffer (2% BSA/RNase-free PBS and 0.2 U/μl Protector RNase inhibitor (Roche)) and re-filtered through a 40 μm strainer. Then, the nuclei suspension was labelled with 10µg/ml DAPI (Merck) and sorted (12-14,000 nuclei per sample) in a 96 well plate directly into 10X RT master mix (without RT Enzyme C) using a FACSAria™ Fusion Flow Cytometer (BD Biosciences). Then 8.3 µl of RT Enzyme C were added to each well.

### 10X Single nuclei RNA-seq

Samples were loaded into the Chromium controller (10X Genomics) for nuclei partition into Gel Bead-In-Emulsions (GEMs). cDNA sequencing libraries were prepared using the Next GEM Single Cell 3’ Reagent Kits v3.1 (10X Genomics, 1000268), following manufacturer’s instructions. Briefly, after GEM-RT clean up, cDNA was amplified during 12 cycles and cDNA QC and quantification were performed on an Agilent Bioanalyzer High Sensitivity chip (Agilent Technologies). cDNA libraries were indexed by PCR using the PN-1000215 Dual Index Kit TT, Set A plate. Size distribution and concentration of 3’ cDNA libraries were verified on an Agilent Bioanalyzer High Sensitivity chip (Agilent Technologies). Finally, sequencing of cDNA libraries was carried out on an Illumina NovaSeq 6000 using the following sequencing conditions: 28 bp (Read 1) + 8 bp (i7 index) + 0 bp (i5 index) + 89 bp (Read 2), to obtain approximately 20-30.000 reads per nucleus.

### Bioinformatic analysis

Various R packages and software were used for the analysis of the samples. Seurat package (4.1.0) was used for the integration of the samples and unsupervised clustering^24^. Monocle-3 was used or trajectory analysis^68^. JavaGSEA was used for gene set enrichment analysis^69^. CellChat was used to study ligand-receptor characterization for cell-cell communication prediction^49^. Raw and processed sequencing data are available under request to the corresponding author. For other detailed methods please refer to the supporting information.

### Functional enrichment analysis

To reveal the precise biological properties of each cluster and in normal or pathological conditions, we used Metascape (http://metascape.org) to perform enrichment analysis including KEGG Pathway, GO Biological Processes, Reactome Gene Sets, Canonical Pathways, CORUM, WikiPathways and PANTER Pathway. Genes with a log_2_FC >0.5 were analysed for each cluster and condition and all genes in the genome were used as the enrichment background.

### Statistics

We confirmed that data on cell population did not follow normal distribution using Shapiro-Wilk test and therefore used nonparametric studies, specifically Mann-Whitney U test, to identify significant differences in cell population between samples. Comparison in gene expression between two or more groups was performed using Kruskal-Wallis test. The level of significance was set at p value <0.05.

This study includes no data deposited in external repositories.

## Acknowledgements

We thank the Single Cell Genomics Group at the National Center of Genomic Analysis (CNAG, Barcelona, Spain), specially to Holger Hein, Ginevra Caratu and Domenica Marchese for their technical assistance. We also thank Rachel Queen and Adrienne Unsworth for their help in the analysis of the bioinformatic data. This work has been funded by grants from Academy of Medical Sciences Professorship Scheme (APR4/1007) and Medical Research Council (MR/W019086/1) to Jordi Díaz-Manera and a grant from La Marató de TV3 (#202034-10) to Xavier Suárez-Calvet.

## Author Contributions

Xavier Suárez-Calvet: Preparing samples for snRNAseq, running assays, planning experiments, writing manuscript, obtaining funding

Esther Fernández-Simón: Preparing samples for snRNAseq, running assays, writing and commenting on the manuscript

Daniel Natera: Follow-up of patients in clinics, collecting clinical data, writing and commenting on the manuscript

Cristina Jou: processing muscles samples, staining of muscle samples, writing and commenting on the manuscript

Patricia Pinol-Jurado: collecting clinical data, commenting on the manuscript Elisa Villalobos: collecting data, writing and commenting on the manuscript

Carlos Ortez: Follow-up of patients in clinics, collecting clinical data, writing and commenting on the manuscript

Alexandra Monceau: Preparing samples for snRNAseq, writing and commenting on the manuscript, Marianela Schiava: collecting data, statistics studies, writing the manuscript

José Verdu-Díaz: bioinformatics analysis, preparing figures for the paper

James Clark: preparing figures for the paper, collecting data, writing the manuscript Zoe Laidler: preparing figures for the paper, collecting data, writing the manuscript Priyanka Mehra: preparing figures for the paper, collecting data, writing the manuscript Rasya Gokul-Nath: bioinformatics analysis, preparing figures for the paper

Jorge Alonso-Perez: Follow-up of patients in clinics, collecting clinical data, writing and commenting on the manuscript

Chiara Marini-Bettolo: Follow-up of patients in clinics, collecting clinical data, writing and commenting on the manuscript

Giorgio Tasca: Follow-up of patients in clinics, collecting clinical data, writing and commenting on the manuscript

Volker Straub: Follow-up of patients in clinics, collecting clinical data, writing and commenting on the manuscript

Michela Guglieri: Follow-up of patients in clinics, collecting clinical data, writing and commenting on the manuscript

Andrés Nascimento: Follow-up of patients in clinics, collecting clinical data, writing and commenting on the manuscript

Jordi Diaz-Manera: planning experiments, obtaining funding, bioinformatics assay, writing and commenting the manuscript

## Supplemental figure legends

**Supplemental Figure 1: Muscle biopsies included in the study.** Representative areas of the muscle biopsies included in the study stained using hematoxilin-eosin and fetal myosin heavy chain antibody.

**Supplemental Figure 2:** UMAP visualization of all nuclei from control and DMD individuals coloured by cluster number before identities were obtained based on the expression of canonical markers.

**Supplemental Figure 3: Heatmap showing the 10 most upregulated genes expressed on each cell cluster.**

**Supplemental Figure 4: Feature-plot showing expression of specific genes in clusters of myonuclei and satellite cells.**

**Supplemental Figure 5: Pseudotime trajectories of satellite cells, regenerative and mature nuclei.**

A) UMAP showing trajectory of satellite cells, regenerative and mature nuclei of control and DMD samples. B) Heatmap showing the expression of genes during pseudotrajectories.

**Supplemental Figure 6: Expression of genes related with myogenesis and cell hypertrophy.** A) Violin plot showing the expression of selected genes belonging to specific pathways involved in myogenic program and cell hypertrophy in myonuclei of fast and slow myofibers, regenerative fibers and satellite cells. B) Graph showing selected molecular pathways involved in the myogenic program and cell hypertrophy in skeletal muscle.

**Supplemental Figure 7:** Expression of genes coding for components of the extracellular matrix by cell type.

Table showing changes in the expression of genes coding for components of the extracellular matrix in control and DMD samples: upregulated in DMD (>0.8 fold increase), not changed (between 0.8 and -0.8 fold increase) and upregulated in controls (>0.8 fold increase).

**Supplemental Figure 8: Gene expression in adipocytes**. A) Volin plot identifying specific genes distinguishing FAPs from adipocytes. B) Molecular pathways upregulated in adipocytes present in muscle samples.

